# Volume and intramuscular fat content of upper extremity muscles in individuals with chronic hemiparetic stroke

**DOI:** 10.1101/687699

**Authors:** Lindsay R. P. Garmirian, Ana Maria Acosta, Ryan Schmid, Jules P. A. Dewald

## Abstract

Stroke survivors often experience upper extremity deficits that make activities of daily living (ADLs) like dressing, cooking and bathing difficult or impossible. Survivors experience paresis, the inability to efficiently and fully activate muscles, which combined with decreased use of the upper extremity, will lead to muscle atrophy and potentially an increase in intramuscular fat. Muscle atrophy has been linked to weakness post stroke and is an important contributor to upper extremity deficits. However, the extent of upper extremity atrophy post hemiparetic stroke is unknown and a better understanding of these changes is needed to inform the direction of intervention-based research. In this study, the volume of contractile tissue and intramuscular fat in the elbow and wrist flexors and extensors were quantified in the paretic and non-paretic upper limb using MRI and the Dixon technique for the first time. Total muscle volume (p≤0.0005) and contractile element volume (p≤0.0005) were significantly smaller in the paretic upper extremity, for all muscle groups studied. The average percent difference between limbs and across participants was 21.3% for muscle volume and 22.9% for contractile element volume. We also found that while the percent intramuscular fat was greater in the paretic limb compared to the non-paretic (p≤0.0005), however, the volume of intramuscular fat was not significantly different between upper limbs (p=0.231). The average volumes of intramuscular fat for the elbow flexors/extensors and wrist flexors/extensors were 28.1, 28.8 and 19.9, 8.8 cm^3^ in the paretic limb and 29.6, 27.7 and 19.7, 8.8 cm^3^ in the non-paretic limb. In short, these findings indicate a decrease in muscle volume and not an increase in intramuscular fat, which will contribute to the reduction in strength in the paretic upper limb.

## Introduction

Stroke is the leading cause of serious long-term disability in the United States with approximately 795,000 new or recurrent strokes occur every year (Writing Group Members et al., 2016). Seventy percent of stroke survivors experience long-term deficits in their upper extremity (Faria-Fortini, Michaelsen, Cassiano, & Teixeira-Salmela, 2011) including difficulty with activities of daily living (ADLs) and tasks that involve reaching. These deficits are attributed to a loss of corticofugal projections that occur post stroke and result in paresis, loss of independent joint control and hypertonicity (Dewald & Beer, 2001; Miller & Dewald, 2012). Associated with neural deficits and resulting disuse, changes in musculoskeletal properties (sarcomere length, fascicle length, stiffness) may also have a negative impact on function, with only a few studies focusing on characterizing these musculoskeletal changes (Jakubowski, Terman, Santana, & Lee, 2017; Landin, Hagenfeldt, Saltin, & Wahren, 1977; Zhao, Ren, Roth, Harvey, & Zhang, 2015). The inability to efficiently and fully activate muscles, combined with decreased use of the upper extremity, will lead to muscle atrophy defined here as a decrease in muscle contractile element volume, and potentially an increase in intramuscular fat. Muscle atrophy, along with other factors will cause unilateral weakness or hemiparesis post unilateral stroke and may contribute to a decline in function. A better knowledge of the level of muscle atrophy in the paretic arm will inform the direction of intervention-based research and assist in improving outcomes for stroke survivors. Therefore, the goal of this study was to determine the amount of atrophy and changes in intramuscular fat that occurs post stroke in muscles of the upper extremity.

Researchers have used a variety of imaging modalities including magnetic resonance imaging (MRI), computed tomography (CT), dual energy x-ray absorptiometry (DEXA) and ultrasound to examine changes in muscle volume in the paretic lower extremity after stroke. The degree of atrophy reported in the lower extremity is variable. Using MRI, Klein et al. found a 24% decrease in the paretic gastrocnemii volume compared to the non-paretic, but no significant difference between limbs for the soleus, deep plantar flexors and peronei (Klein, Brooks, Richardson, McIlroy, & Bayley, 2010). Ramsay et al. also used MRI and reported a decrease in volume between 1% and 33% for fourteen muscles in the paretic lower extremity compared to the non-paretic and a 14% increase in the paretic gracilis (Ramsay, Barrance, Buchanan, & Higginson, 2011).

Fewer studies exist examining atrophy in the upper extremity, due to technical challenges that make imaging the upper extremity more difficult compared to the lower extremity. These technical difficulties include wrap around artifacts due to the close proximity to the trunk as well as decreased field strength at the edge of the bore (field inhomogeneity), where the upper-extremity is most comfortably scanned. A review paper by English et al. in 2010 reported six studies that looked at atrophy of the upper extremity in chronic hemiparetic stroke. These six studies reported a range of values from no significant atrophy up to a 25% decrease in volume (English, McLennan, Thoirs, Coates, & Bernhardt, 2010). Four of these studies used DEXA and reported total upper limb lean tissue mass and one study used ultrasound and reported muscle thickness. The sixth study was a pilot study with six subjects, done in 2006 that used MRI and found a 25% decrease in triceps cross-sectional area and no significant difference in biceps cross-sectional area (Ploutz-Snyder, Clark, Logan, & Turk, 2006). In 2012, Triandafilou et al. reported a 15% difference in cross-sectional area for muscles controlling the index finger using ultrasound (Triandafilou & Kamper, 2012). This study looks to improve upon these results by increasing the number of participants studied, using MRI, the gold standard imaging modality to assess muscle atrophy(Fuller et al., 1999), and examining muscle volume which is a more robust measure than cross-sectional area or thickness.

This study also paints a more complete picture of the muscular changes that occur post stroke, by measuring and accounting for changes in intramuscular fat using the Dixon technique (Dixon, 1984). Studies have shown that MRI can be used to accurately quantify intramuscular fat in vivo, as compared to muscle biopsy (A. C. Smith et al., 2014). An increase in the percentage of intramuscular fat has been reported in a variety of pathologies including patients with chronic whiplash associated disorder(Elliott et al., 2010; 2006), after rotator cuff tear (Fuchs, Weishaupt, Zanetti, Hodler, & Gerber, 1999) and in individuals with spinal cord injury (Elder, Apple, Bickel, Meyer, & Dudley, 2004). Intramuscular fat has not been studied in the upper extremity post stroke; however, studies have examined changes in the lower extremity. Ramsay et al. reported a significantly greater percent intramuscular fat in ten of fifteen paretic lower extremity muscles compared to the non-paretic (Ramsay et al., 2011).

Historically, changes in intramuscular fat have been reported only as percentages, the volume of intramuscular fat normalized by total muscle volume. However, this method does not provide insight into whether intramuscular fat volumes have increased, muscle tissue volumes have decreased or both. The non-paretic limb provides a convenient within subject control, that allows us to compare the volume of intramuscular fat between limbs and infer whether the volume of intramuscular fat has changed post stroke.

In short, the purpose of the present study was to quantify the amount of contractile tissue and intramuscular fat in upper extremity muscles in the paretic and non-paretic limb of chronic hemiparetic stroke individuals for the first time. We hypothesized that the volume of contractile tissue (total muscle volume minus intramuscular fat) would be decreased in the paretic limb, compared to the non-paretic limb.

## Methods

### Participants

The Rehabilitation Institute of Chicago’s (currently known as the Shirley Ryan AbilityLab) clinical neuroscience research registry database was used to recruit ten individuals with chronic hemiparetic stroke (7 males, 3 females; mean ± SD age: 59.2 ± 2.6 years, Table 3.1). All lesions had occurred at least 3 years prior (mean ± SD: 11.0 ± 2.5 years, Table 3.1) Seven participants had right-sided hemiparesis and three had left-sided hemiparesis. Participants were excluded if they had a severe concurrent medical problem, an acute or chronic painful or inflammatory condition of either upper extremity or diabetes. All participants gave informed consent for participation in the study, which was approved by the institutional Review Board of Northwestern University. All experimental procedures were conducted according to the Declaration of Helsinki.

**Table 3.1.** Participant demographics

| Participant | Gender | Group | Affected | Dominant | Age | Years Post | Fugl-Meyer |
| --- | --- | --- | --- | --- | --- | --- | --- |
|  |  |  |  | limb | (years) | stroke | Upper-Extremity |
| 1 | M | Stroke | R | R | 55.8 | 3.7 | 24 |
| 2 | F | Stroke | R | R | 63.1 | 8.0 | 30 |
| 3 | M | Stroke | L | R | 45.6 | 5.4 | 33 |
| 4 | F | Stroke | R | R | 60.4 | 12.8 | 25 |
| 5 | M | Stroke | R | R | 65.4 | 8.5 | 18 |
| 6 | F | Stroke | L | R | 61.6 | 28.1 | 16 |
| 7 | M | Stroke | R | R | 69.4 | 19.0 | 12 |
| 8 | M | Stroke | R | R | 63.5 | 14.7 | 48 |
| 9 | M | Stroke | L | R | 62.2 | 5.0 | 47 |
| 10 | M | Stroke | R | R | 44.6 | 5.0 | 38 |
| **Mean (SD)** |  |  |  |  | **59.2 (2.6)** | **11.0 (2.5)** | **9.1 (4.0)** |

### Fugl-Meyer Assessment

All participants were assessed using the Fugl-Meyer Assessment (FMA), a stroke specific, performance-based motor impairment tool. The FMA measures reflex activity, active movment of the shoulder, elbow and hand, loss of independent joint control and coordination. All participants had moderate to severe upper limb motor impairment according to the Fugl-Meyer Motor Assessment upper extremity (FMA-UE) portion (mean ± SD: 22.7 ± 7.2, Table 3.1).

### Magnetic Resonance Imaging

A 3-D dual-echo gradient, fat-water separation MRI technique (two-point Dixon) was used to acquire the data (repetition time = 7ms, echo time 1 = 2.45ms, echo time 2 = 3.675ms, flip angle =12°, imaging matrix = 256 × 304, slice thickness = 3mm). Two echo times were used to estimate percent fat, one when water and fat are in phase (2.45ms) and one when water and fat are out of phase (3.675ms). Two additional sets of images were derived from the original images, one in which water was the main contributor to signal intensity (water-only images) and one in which fat was the main contributor to signal intensity (fat-only images). A standard 12-channel upper extremity receiver coil was used to improve the signal-to-noise ratio.

Participants were asked to lay supine outside the MRI bore and their elbow and wrist were ranged for approximately five minutes, or until their elbow and wrist achieved a neutral position at rest. A forearm-hand orthosis was used to keep the participant’s wrist and fingers in a neutral position and the upper extremity was fixed at the participant’s side using cloth straps and foam supports. Once inside the bore, a localizer scan was performed followed by two overlapping scans of the upper extremity. Two scans were performed to maximize coverage area, resolution and signal to noise ratio. One scan covered the upper-extremity from the acromion to the head of the radius while the second scan spanned from the medial epicondyle to the proximal row of carpal bones. Visual inspection of the acquired images was performed immediately following data acquisition to guarantee the absence of artifacts. If artifacts were observed, the scans were repeated. This process was repeated for both the paretic and non-paretic upper limbs for all ten participants.

### Segmentation

The MR data was segmented using Analyze software (Analyze 12.0, Analyze Direct, Overland Park, KS). The boundaries of the muscles of interest were manually segmented within the epimysium bilaterally, using the axial images from both scans. For each set of images, overlapping segmentation areas were removed by exporting the images of the reconstructed muscles from the two scans to Matlab, aligning the images using the medial and lateral epicondyles and the olecranon and removing redundant pixels.

Analyze software was used to calculate muscle volume as well as percent intramuscular fat. The percent intramuscular fat was determined from a ratio of the intensity of the signal from the fat-only images normalized to the sum of the intensity from the water-only images and fat-only images. The volume of intramuscular fat was determined by multiplying the total muscle volume by the percent intramuscular fat. Contractile element volume was then determined for each muscle by subtracting the volume of intramuscular fat from the total muscle volume. The interlimb difference in volume was calculated as a proxy measure for atrophy and was defined as the relative difference in contractile element volume between paretic and non-paretic limbs, normalized by the contractile element volume of the non-paretic limb.

A single rater segmented all of the muscles of interest once and repeated segmentations for one participant on a separate day, in order to calculate intra-rater reliability. Additionally, a second rater segmented the muscles for one participant, to establish inter-rater reliability of the method.

Individual muscles were grouped into functional groups (elbow flexors/extensors and wrist flexors/extensors) according to peak moment arm in the following position: 15° shoulder flexion, 30° shoulder abduction, 90° elbow flexion and in neutral with respect to pronation/supination and wrist and finger posture (Table 3.2) (Holzbaur, Delp, Gold, & Murray, 2007; Holzbaur, Murray, & Delp, 2005).

**Table 3.2.** Functional groups of muscles

| **Elbow Flexors** | **Elbow Extensors** |
| --- | --- |
| Biceps brachii (BIC) | Triceps brachii (TRI) |
| Brachialis (BRA) | Anconeus (ANC) |
| Brachioradialis (BRD) | Supinator (SUP) |
| Pronator teres (PT) |  |

| **Wrist Flexors** | **Wrist Extensors** |
| --- | --- |
| Flexor carpi radialis (FCR) | Extensor carpi radialis longus (ECRL) |
| Flexor carpi ulnaris (FCU) | Extensor carpi radialis brevis (ECRB) |
| Palmaris longus (PL) | Extensor carpi ulnaris (ECU) |
| Flexor digitorum superficialis (FDS) | Extensor digitorum communis (EDC) |
| Flexor digitorum profundus (FDP) | Extensor digit minimi (EDM) |
| Flexor pollicis longus (FPL) | Extensor indicis proprio (EIP) |
| Abductor pollicis longus (APL) | Extensor pollicis longus (EPL) |
|  | Extensor pollicis brevis (EPB) |

### Statistics

Pearson correlation coefficients were calculated to determine the intra-rater and inter-rater reliability for contractile element volume and intramuscular fat volume for both the paretic and non-paretic limbs. Four repeated measures analyses of variance (ANOVAs) were used to determine the effect of limb (paretic, non-paretic) on the dependent variables muscle volume, percent intramuscular fat, contractile element volume and intramuscular fat volume. Pearson correlation coefficients were used to determine if there was a significant correlation between percent difference in contractile element volume for the paretic limb compared to the non-paretic and impairment level as measured by the FMA arm subscore. A p-value of ≤0.05 was considered significant.

## Results

### Intra-rater and Inter-rater Reliability

Intra-rater reliability was high for both contractile element volume and intramuscular fat volume for both the paretic and non-paretic limbs (paretic contractile element volume correlation coefficient, = 0.98, non-paretic contractile element volume correlation coefficient= 0.97, paretic intramuscular fat volume correlation coefficient= 0.97, non-paretic intramuscular fat volume correlation coefficient= 0.96). Additionally, inter-rater reliability was also high for both outcome measures (paretic contractile element volume correlation coefficient= 0.96, non-paretic contractile element volume correlation coefficient, = 0.97, intramuscular fat volume paretic correlation coefficient= 0.96, non-paretic intramuscular fat volume correlation coefficient= 0.95).

### Percent Intramuscular Fat

The percent intramuscular fat was significantly greater in the paretic compared to the non-paretic limb, across participants and muscle groups (p≤0.0005), as illustrated in Figure 3.1. For all four muscle groups, the average percent intramuscular fat was significantly greater in the paretic limb compared to the non-paretic limb across participants (Fig 3.1). The average percent intramuscular fat for the paretic limb was 8.7%, 8.9% and 8.9%, 8.8% for the elbow flexors/extensors and wrist flexors/extensors respectively, and 7.2%, 6.5% and 6.7%, 6.7% for the non-paretic limb (Table 3.3).

**Figure 3.1.**
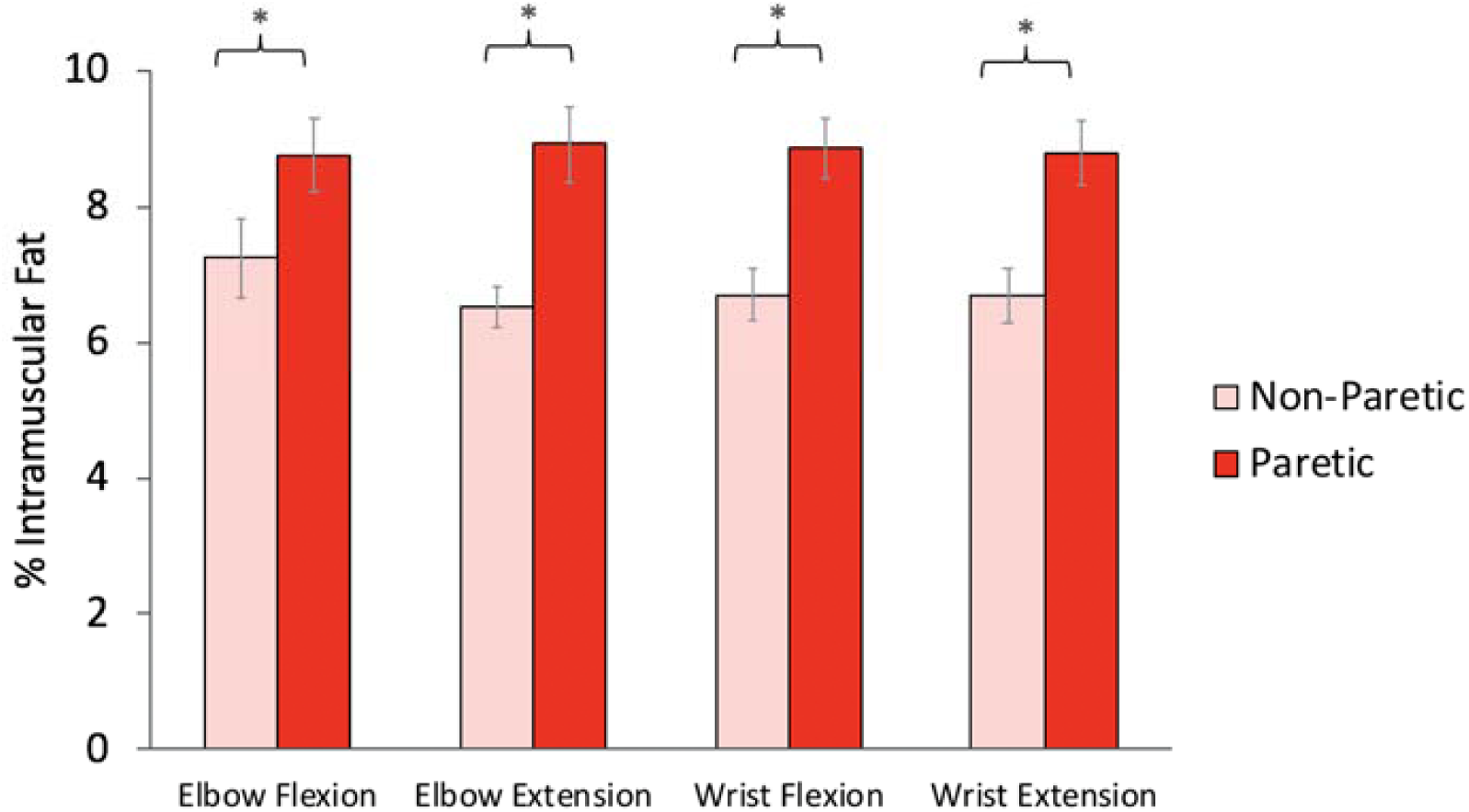
Percent intramuscular fat for the elbow flexors/extensors and wrist flexors/extensors of the paretic and non-paretic limbs. Results are the average for ten participants ± one standard error.

**Table 3.3.** Mean (standard error) percent intramuscular fat, contractile element volume, muscle volume and intramuscular fat volume for the non-paretic and paretic limbs as well as the interlimb difference in volume for the paretic limb compared to the non-paretic.

|  | Muscle group | Non-Paretic | Paretic | Interlimb difference (%) |
| --- | --- | --- | --- | --- |
| Percent Intramuscular Fat | Elbow Flexors* | 7.2 (0.59) | 8.7 (0.52) |  |
|  | Elbow Extensors* | 6.5 (0.30) | 8.9 (0.55) |  |
|  | Wrist Flexors* | 6.7 (0.38) | 8.9 (0.44) |  |
|  | Wrist Extensors* | 6.7 (0.40) | 8.8 (0.45) |  |
| Contractile Element Volume (cm<sup>3</sup>) | Elbow Flexors* | 377.5 (45.4) | 301.1 (39.3) | 19.7 (5.3) |
|  | Elbow Extensors* | 394.1 (39.8) | 297.2(40.6) | 25.7 (5.1) |
|  | Wrist Flexors* | 271.2 (28.9) | 207.1(21.1) | 21.5 (4.9) |
|  | Wrist Extensors* | 122.4 (13.5) | 93.2 (13.9) | 25.3 (5.1) |
| Muscle Volume (cm<sup>3</sup>) | Elbow Flexors* | 407.1 (49.9) | 329.2(42.6) | 18.4 (5.33)) |
|  | Elbow Extensors* | 421.8 (43.1) | 326.1 (44.2) | 23.9 (5.07) |
|  | Wrist Flexors* | 290.9 (31.5) | 227.0 (23.0) | 19.7 (5.08) |
|  | Wrist Extensors* | 131.2 (14.7) | 102.1 (15.2) | 23.7 (5.06) |
| Intramuscular Fat Volume (cm<sup>3</sup>) | Elbow Flexors | 29.6 (5.4) | 28.1 (3.8) | 0.37(6.77) |
|  | Elbow Extensors | 27.7(3.7) | 28.8 (4.1) | -4.2 (8.7) |
|  | Wrist Flexors | 19.7 (3.0) | 19.9 (2.2) | -7.2(8.3) |
|  | Wrist Extensors | 8.8 (1.3) | 8.8 (1.3) | -0.46 (7.0) |

### Muscle Volume

Muscle volume (the sum of contractile element and intramuscular fat) was significantly reduced in the paretic limb compared to the non-paretic limb across participants, for all muscles (p≤0.0005), as illustrated in Figure 3.2 (sum of grey and white bars). The average percent difference between limbs and across participants was 18.4%, 23.9% and 19.7%, 23.7% for the elbow flexors/extensors and wrist flexors/extensors, respectively (Table 3.3).

**Figure 3.2.**
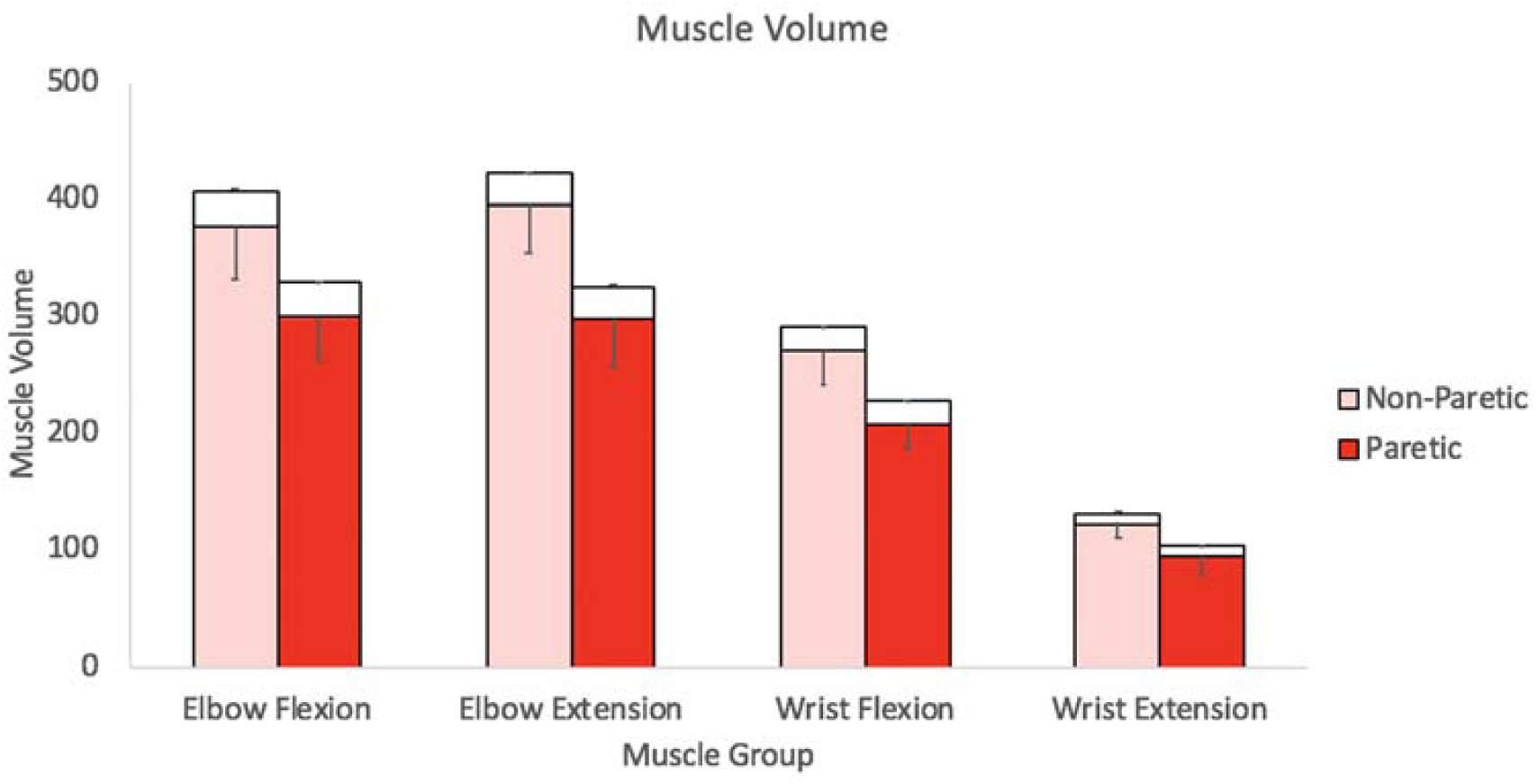
Total muscle volume displayed as intramuscular fat (white bars) volume stacked on top of contractile element volume (colored bars) for the elbow flexors/extensors and wrist flexors/extensors of the non-paretic (light red) and paretic (dark red) limbs. Results are the average for ten participants plus one standard error (intramuscular fat) and minus one standard error (contractile element).

### Contractile Element Volume

The contractile element volume was significantly reduced in the paretic compared to the non-paretic limb across muscles (p≤0.0005), as illustrated in Table 3.3. The percent difference between limbs was 19.7%, 25.7%, and 21.5%, 25.3% for the elbow flexors/extensors, wrist flexors/extensors, respectively (Table 3.3).

### Intramuscular Fat Volume

No significant differences in intramuscular fat volume were found for the paretic limb compared to the non-paretic limb (p=0.231) across participants, as illustrated in Figure 3.2 and Table 3.3.

### Impairment level versus contractile element percent difference between limbs

There was no statistically significant correlation between each participant’s impairment level, as measured by the FMA and the contractile element volume interlimb difference for the elbow flexors (*R*^*2*^ = 0.29 *P* = .106), wrist flexors (*R*^*2*^ = 0. 1378 *P* = .291) or wrist extensors (*R*^*2*^ = 0. 052, *P* = .395). Interestingly, the correlation between the FMA and contractile element interlimb difference for the elbow extensors (*R*^*2*^ = 0.4605, *P* = .031).

## Discussion

This study presents percent intramuscular fat, muscle volume, contractile element volume and intramuscular fat volume of the paretic and non-paretic elbow flexors/extensors and wrist flexors/extensors in ten individuals with chronic hemiparetic stroke. This is the first time that a comprehensive characterization of volume is performed in muscles of the upper extremity post stroke based on MR imaging and the Dixon technique. We found that total muscle volume and contractile element volume were significantly smaller in the paretic upper extremity, for all muscle groups studied. We also found that while the percent intramuscular fat was greater in the paretic limb compared to the non-paretic, however the volume of intramuscular fat was not significantly different between limbs.

Results from previous studies examining the effect of stroke on muscle atrophy in the paretic arm vary from no significant effect up to 25% atrophy. The results from this study agree with five of six previous studies that show significant atrophy does occur (Iversen, Hassager, & Christiansen, 1989; Pang & Eng, 2005; Ploutz-Snyder et al., 2006) (Carin-Levy et al., 2006; Ryan, Dobrovolny, Smith, Silver, & Macko, 2002; Triandafilou & Kamper, 2012).Our results demonstrate an average percent difference between the paretic and non-paretic limb of 19.7% in the elbow flexors up to a 25.7% difference in the elbow extensors, which is greater than past studies that report a range between 7% and 25%. This disparity may be due to the fact our participants, on average, were more impaired than compared to past studies. Many of those studies consisted of convenience samples (Carin-Levy et al., 2006; Ryan, Dobrovolny, Silver, Smith, & Macko, 2000) and subjects were likely less impaired than our participants. Additionaly, our participants were 12.7 years post stroke and some of the past studies examined participants with less chronicity, including less than one year post stroke (Carin-Levy et al., 2006; Iversen et al., 1989; Moukas et al., 2002). Additionally, past studies of the upper extremity used DEXA or measured cross-sectional area with ultrasound or MRI, methods that do not account for intramuscular fat and could result in errors due measurement of a 3-dimensional structure with a 2-dimensional imaging technique. Our results agree with past MRI studies in the lower extremity that used muscle volume as an outcome measure and reported up to a 33% difference in muscle volume between lower extremities (Ramsay et al., 2011).

This paper is the first to examine intramuscular fat using MRI in the upper extremity post stroke. Ramsay et al. reported an increase in the percent intramuscular fat in the paretic lower extremity compared to the non-paretic using a T1 intensity threshold (Ramsay et al., 2011). Iverson and Jorgensen have both reported an increase in total limb fat mass (intramuscular and subcutaneous fat combined) in the paretic limb compared to the non-paretic (Iversen et al., 1989; Jørgensen & Jacobsen, 2001). Results obtained using the Dixon technique to evaluate the intramuscular fat content of individual muscles, show that the percent intramuscular fat was two to three percentage points greater in the paretic limb compared to the non-paretic. However, when comparing the volume of intramuscular fat between arms, no significant differences were found.

These results demonstrate the importance of reporting the volume of intramuscular fat as opposed to reporting fat as a percentage of total muscle volume. The paretic muscle may appear fattier than the non-paretic when examining the percent fat values however, the volumetric measurements show that this is due to a decrease in total muscle volume, not an increase in intramuscular fat volume. Past studies that have only examined fat as a percentage of total muscle volume are therefore inconclusive and cannot be used to determine if the observed increase in percent intramuscular fat is due to reductions in total muscle volume or an increase in intramuscular fat volume. This paper suggests that standard practice regarding reporting of intramuscular fat should include volume data in addition to percentages. While not all pathologic populations have a within participant control to compare volumes, an alternative to avoid this issue is to track changes in fat and volume across time.

This paper highlights the importance of studying changes in musculoskeletal properties after hemiparetic stroke, in addition to the neurological deficits that have been studied previously. For decades, stroke research has focused on the neurological deficits that occur including inability to activate motor units (Klein et al., 2010), reduced mean firing rates of motor units (Miller et al., 2014; Rosenfalck & Andreassen, 1980; Tang & Rymer, 1981) as well as possible loss of motor units (McComas, Sica, Upton, & Aguilera, 1973). However, researchers are just beginning to understand changes in musculoskeletal properties including atrophy (Klein et al., 2010; Ramsay et al., 2011), force-length, force-velocity, fiber type changes (Hachisuka, Umezu, & Ogata, 1997), passive stiffness and fascicle length (Li, Tong, & Hu, 2007; Zhao et al., 2015) all key for characterizing the force producing properties of muscle. Deficits in muscle volume will be proportional to deficits in strength assuming that pennation angle, optimal fiber length and specific tension remain unchanged following stroke. Our findings indicate a 19.7-25.7% (Table 3.3) decrease in contractile element volume, which will be associated with a reduction in strength and may be related to the impairment level of chronic stroke survivors. Our findings show a correlation between impairement level and elbow extensor atrophy, suggesting that greater impairment may result in greater disuse and thus greater atrophy. Interestingly, this trend was not found for the elbow flexors, wrist flexors or wrist extensors, only the elbow extensors, a muscle group that may be activated less often in inviduals with severe impairment due to expression of the flexion synergy(Miller McPherson & Dewald, 2019).

There are several limitations of this study. The sample size is small due to the amount of time required to manually segment the muscles of interest. Additionally, imaging the upper extremity requires the participant to be able to lay still with their arm at the center of the MRI bore (where field strength is greatest) for an extended period of time. In most populations, this position is usually attained by having subjects lay prone with their arm extended above their head. This position is not feasible due to the passive range of motion restrictions common in individual with chronic stroke. Therefore, we scanned participants in a supine position with their arm at their side with the trunk at the edge and the arm in the center of the bore. The limited sample size, paired with the fact that only moderately and severely impaired participants were included, may be the reason there was only a trend in the correlation between impairment and atrophy of the triceps.

This is the first study, to our knowledge, that has determined the volume of muscle and intramuscular fat in the upper extremity in participants with chronic hemiparetic stroke. Our study confirms that there is a significant reduction in muscle volume between the non-paretic and paretic and limb in the muscle groups studied but that there is no significant difference in the volume of intramuscular fat between limbs. Our study shows that muscle atrophy may be related to upper extremity impairment in chronic stroke survivors. Finally, since changes in the percentage of intramuscular fat were shown to be due to differences in contractile element volume and not intramuscular fat volume, this study illustrates the importance of reporting differences in intramuscular fat as volumes and not solely percentages.

